# Sequence-Directed Covalent Protein-RNA Linkages in a Single Step Using Engineered HUH-Tags

**DOI:** 10.1101/2024.08.13.607811

**Authors:** Adam T. Smiley, Natalia Babilonia-Díaz, August J. Krueger, Hideki Aihara, Kassidy J. Tompkins, Andrew C.D. Lemmex, Wendy R. Gordon

## Abstract

Replication-initiating HUH-endonucleases (Reps) are enzymes that form covalent bonds with single-stranded DNA (ssDNA) in a sequence specific manner to initiate rolling circle replication. These nucleases have been co-opted for use in biotechnology as sequence specific protein-ssDNA bioconjugation fusion partners dubbed ‘HUH-tags’. Here, we describe the engineering and *in vitro* characterization of a series of laboratory evolved HUH-tags capable of forming robust sequence-directed covalent bonds with unmodified RNA substrates. We show that promiscuous Rep-RNA interaction can be enhanced through directed evolution from nearly undetectable levels in wildtype enzymes to robust reactivity in final engineered iterations. Taken together, these engineered HUH-tags represent a promising platform for enabling site-specific protein-RNA covalent bioconjugation in vitro, potentially mediating a host of new applications and offering a valuable addition to the HUH-tag repertoire.

## Introduction

Synthetic protein-RNA interaction has proven itself to be an indispensable utility in basic science and biotechnology alike. One prominent example is the MS2 bacteriophage coat protein, which binds a cognate RNA hairpin with high specificity and affinity^1^. This system has been widely deployed in a diversity of applications, ranging from Cas9-mediated genome visualization^2^, to the induction of trans-splicing in coordination with Cas13^3^, and even in the design of nanocages to protect RNAs in synthetic cell-to-cell mRNA delivery systems^4^. While affinity-based protein-RNA interactions have been employed extensively, covalent linkage between proteins and RNA offers distinct advantages, such as irreversible binding and enhanced stability through direct linkage of an exposed end, protecting the molecule from exonuclease degradation. However, covalent linkage also presents unique challenges, as it often requires synthetic methods that rely on the chemical modification of surface exposed residues in order to facilitate the formation of a stable protein-RNA bond, which could inhibit the labeled protein’s biological function^5-8^.

Enzymatic bioconjugation techniques have the potential to mitigate the negative aspects of synthetic methods by providing a more straightforward and efficient means by which to achieve covalent linkage between proteins of interest and RNA sequences of interest. The SNAP-^9^, CLIP-^10^, and HALO-tag^11^ systems have emerged as a highly versatile suite of bioconjugation tags that can agnostically append to a molecular substrate so long as it contains their specific chemical moiety, making them promising candidates for mediating protein-RNA bioconjugation. Indeed, SNAP-tags have already been successfully deployed in this context. For example, these bioconjugation tags have been directly fused to adenosine deaminases acting on RNA (ADAR) enzymes to mediate covalent fixation of a guiding RNA and enable temporally controlled site-specific RNA editing^12^. Further, SNAP-tags have also been employed in a generalizable strategy to mediate designer protein-RNA covalent interaction more broadly^13^. While these examples demonstrate the potential of enzymatic bioconjugation for protein-RNA linkage, the expensive chemical moieties required to enable SNAP-tag based strategies somewhat limits their broad adoption in these applications. This limitation has prompted the exploration of alternative enzymatic strategies that could enable more cost-effective and accessible protein-RNA bioconjugation. One promising avenue lies in the repurposing of naturally occurring enzymes that form covalent bonds with nucleic acids as part of their native function.

The HUH-endonuclease superfamily is a diverse group of ssDNA-interacting sequence-specific nucleases that contain an eponymous motif consisting of a pair of conserved metal-coordinating histidine (H) separated by a bulky hydrophobic residue (U)^14,15^. This superfamily is comprised of replication initiator proteins involved in rolling-circle replication, relaxases involved in plasmid conjugation, and a diversity of DNA transposases ranging from the prokaryotic insertion sequence transposase families to the eukaryotic Helitron rolling-circle transposases and the newly discovered Replitron transposases^14-21^. Every member of this superfamily employs a catalytic tyrosine to cleave and then covalently link to ssDNA via a 5’ phosphotyrosine linkage.

These nucleases have been co-opted for application in biotechnology as functional fusion partners (HUH-tags) that enable facile and robust sequence-directed protein-ssDNA bioconjugation^20,22^ (Figure 1A). Indeed, HUH-tags have been applied in CRISPR-based genome engineering as a means by which to covalently tether a repair template to Cas9 in order to enhance homology directed repair for gene editing and knock-in^23-25^, and in receptor-specific cell targeting of adeno-associated viruses with capsid-fused HUH-tags linked to ssDNA-conjugated guiding antibodies to control viral tropism^26,27^. Moreover, HUH-tag utilities extend beyond genome engineering into a diversity of applications, ranging from DNA-coordinated enzyme assembly^28^ to high-throughput DNA tension gauge tether-based tension profiling^29^. HUH-tags are amenable to these applications due to their modest size and their ability to rapidly and robustly form non-labile sequence-directed covalent linkages with unmodified ssDNA.

**Figure 1:**
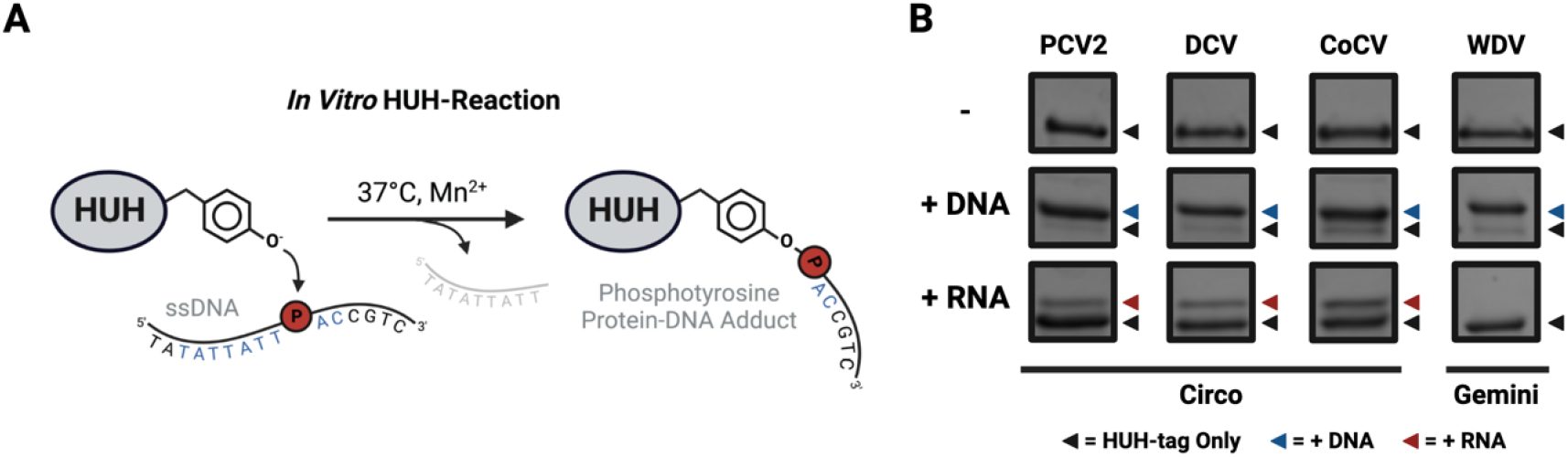
Promiscuous interaction with RNA substrates. (A) Graphical overview of an in vitro HUH-tag bioconjugation reaction. (B) in vitro HUH-tag bioconjugation reactions visualized via SDS-PAGE with either DNA (+ DNA) or RNA (+ RNA) substrates harboring an HUH-tag cleavage motif with reps derived from either *Circoviridae* (Circo), or *Geminiviridae* (Gemini). Reactions were performed in final concentrations of 3 μM HUH-tag 30 μM of indicated oligo in 50 mM HEPES pH 8.0, 50 mM NaCl, 1 mM DTT, and 1 mM MnCl_2_ for 24 hours at 37°C.

We have discovered that a minority of viral Rep HUH-endonucleases from the *Circoviridae* family are able to form promiscuous sequence-directed covalent bonds to RNA, but with greatly reduced efficiency in comparison to cognate reactions with ssDNA substrates. Here, we describe the engineering and *in vitro* characterization of a series of engineered RNA-interacting HUH-tags derived from the HUH-endonuclease domain of the replication protein from Porcine Circovirus 2 (PCV2).

## Results

### Identifying Promiscuous Protein-RNA Covalent Linkage

In our previous work, we introduced HUH-tags as functional fusion partners capable of mediating sequence-directed protein-ssDNA covalent bioconjugation in a single step^20,22^. HUH-tags derived from both replication initiating HUH-endonucleases (reps), responsible for initiating rolling circle replication in extrachromosomal plasmids and viral genomes, and relaxase HUH-endonucleases, responsible for mediating conjugative plasmid transfer in prokaryotes, have been deployed in a wide diversity of applications^14,15,20,22^. Recently, the reps have emerged as the preferred HUH-tag type due to their modest size, compact cleavage motifs, and robust reactivity^20^. Moreover, our extensive knowledge of their sequence specificity^20^, coupled with the recent development of computational tools^30^ to aid in their substrate design, have further simplified their deployment and increased the utility of rep-based HUH-tags. While reps can be identified in a wide diversity of bacterial, archaeal, and viral species, the HUH-tags are primarily co-options of reps belonging to the ssDNA virus phylum *Cressdnaviricota*^31^ – typically from the *Circoviridae, Geminiviridae*, and *Nanoviridae* families. Interestingly, we recently discovered that reps originating from *Circoviridae* can exhibit some promiscuity which enables interaction with ssRNA, albeit with dramatically lower efficiency in comparison to reactions with cognate ssDNA substrates (Figure 1B).

*In vitro* reactions with a panel of HUH-tags – three derived from the circoviruses PCV2, duck circovirus (DCV) and columbid circovirus (CoCV) and one from the geminivirus wheat dwarf virus (WDV) – analyzed by SDS-PAGE showed that overnight incubation under favorable conditions yielded over 90% protein-ssDNA adduct formation for all four reps tested.

Interestingly, reps derived from Circoviridae, but not Geminiviridae, were also capable of forming sequence-directed covalent linkages with RNA, albeit with much lower efficiency (roughly 15%, 12%, and 20% adduct formation for PCV2, DCV, and CoCV, respectively) compared to reactions with cognate ssDNA substrates.

The discovery of promiscuous activity on RNA substrates in *Circoviridae*-derived reps presents an exciting opportunity to develop RNA-interacting HUH-tags – fusion proteins capable of mediating simple and efficient sequence-directed protein-RNA covalent linkage on unmodified RNA substrates. Enhancing the RNA-interacting capabilities of these enzymes will likely require protein engineering, and encouragingly, we have previously demonstrated that Reps from PCV2 and WDV are amenable to rational engineering^20^, making PCV2 a suitable scaffold to enhance this interaction. While enzymes are typically engineered rationally through site-directed mutagenesis and in vitro screening^32^, high-throughput methods, such as yeast display^33^ or droplet microfluidics^34^, have also been employed successfully across a wide variety of examples. Indeed, yeast display has been used to engineer other bioconjugation-mediating enzymes, such as sortase^35^, a transpeptidase that catalyzes sequence-specific intermolecular protein ligation. Moreover, this method is commonly used to engineer meganucleases for targeted gene editing applications^36^, further highlighting the versatility and effectiveness of yeast display in the directed evolution of a diverse range of enzymes. The self-labeling nature of Reps makes them ideal candidates for yeast display methods, which often rely on fluorescence-based selection using flow cytometry, as engineered variants with enhanced properties can accumulate higher fluorescence in a gain-of-signal (i.e., ‘signal on’) selection scheme.

### Developing a Directed Evolution Approach for Rep Engineering

Inspired by the success of other high throughput enzyme engineering efforts, we developed a yeast surface display platform to enhance Rep-RNA interaction. Yeast surface display is a powerful tool for directed evolution that employs *Saccharomyces cerevisiae* to establish a link between genotype (i.e., enzyme coding sequence) and phenotype (i.e., enzyme function). In this system, mutant enzymes are tethered to the yeast cell surface through plasmid-driven expression as fusions with a cell wall-anchoring protein which facilitates their export and attachment to the cell’s exterior. Because Reps are self-labeling enzymes, they are superb candidates for yeast display-based engineering because the method enables high-performing enzymes to accumulate more functionalized substrate (either biotinylated or fluorescently labeled RNA) on the cell surface. Consequently, cells displaying superior enzymes can be effectively selected and enriched through magnetic-activated cell sorting (MACS) with streptavidin beads or fluorescence-activated cell sorting (FACS) (Figure 2A).

**Figure 2:**
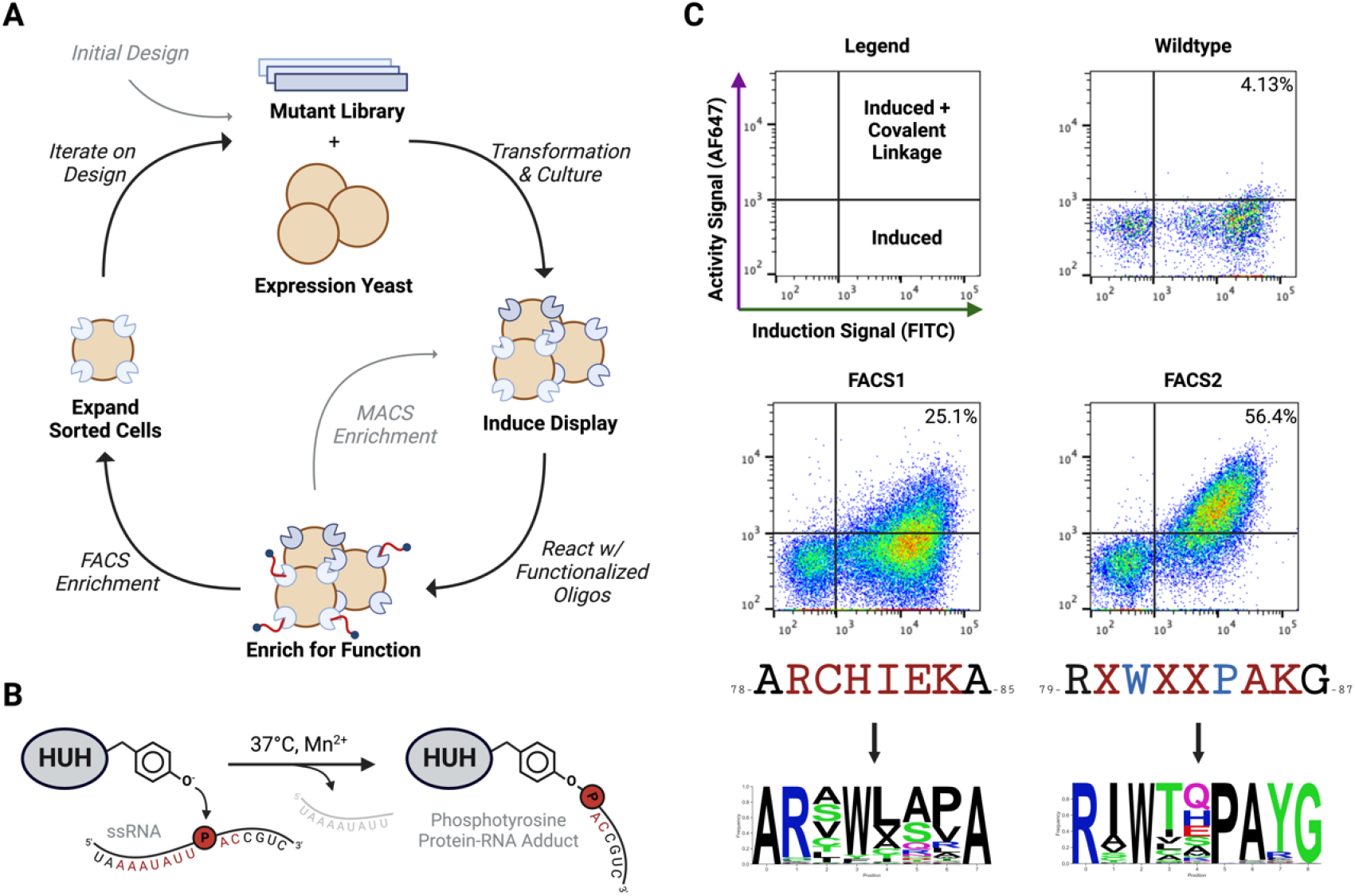
Overview of the directed evolution process. (A) High level overview of the yeast surface display-mediated selection scheme employed by this manuscript. (B) Graphic depiction of the desired engineered RNA HUH-tag reaction. (C) Results of yeast surface display directed evolution. The flow cytometry plots indicate the population of yeast inducing enzyme expression (shifts on the X axis) and covalently linking to fluorescently labeled RNA (shifts on the Y axis) for the wildtype enzyme as well as the heterogenous results of the FACS sorting from the first and second rounds of engineering (FACS1 and FACS2). The residues underneath their associated flow plots indicate the mutagenic library being selected on where red indicates replacement with a degenerate codon, blue indicates convergence to a residue from a previous round of engineering, and X indicates a lack of convergence and a need for re-engineering. Sequence logos of library enrichment post sort are shown below the library designs.

In initial tests, we expressed the wild-type (WT) PCV2 enzyme on the surface of yeast and incubated it with fluorescently labeled RNA substrates in the presence of Mn^2+^ to assess if baseline promiscuous activity is detectable in this context. Flow cytometry analysis revealed that approximately 4% of the total yeast population displayed proteins labeled with fluorescent RNA (Figure 2B & C). However, the magnitude of the fluorescent signal was quite low, indicating that the reaction efficiency of the WT enzyme to RNA was modest, but detectable, which is consistent with our previous in vitro observations (Figure 1B).

While the WT enzyme’s RNA-linking activity is quite low on the surface of yeast, it provides a detectable signal that can serve as a baseline for selecting variants with higher levels of activity. Thus, we designed libraries using site-saturation mutagenesis to screen for enzyme variants with enhanced function using our yeast display system. Specifically, we synthesized mutagenic libraries by using assembly PCR^37^ to replace residues in regions of interest with ‘NNK’ degenerate codons that encode all twenty amino acids and one stop codon. This strategy enables comprehensive screening of all potential combinations of mutations across positions of interest, facilitating an exhaustive search of the local sequence space to identify enzymes with optimal function. We focused on mutating regions of the enzyme implicated in protein-ssDNA interaction in an effort to better accommodate the highly comparable RNA substrates. Recent co-crystal structures of Reps in complex with their substrates revealed the single-stranded DNA bridging motif (sDBM)^20^ – a motif seemingly universal to the HUH-endonuclease superfamily – that is a key regulator of substrate interaction and sequence specificity. In particular, a stretch of six residues (positions 79-84) of this motif are shown to makes extensive contacts with the ssDNA substrate in PCV2. Thus, we targeted this region for our initial round of mutagenesis and screening, transforming the assembled mutagenic library into yeast by leveraging the organism’s natural ability to mediate plasmid recombination^38^.

To select for enzymes with optimized function, we performed two initial rounds of MACS with increasing stringency to enrich the population for highly functional variants followed by a single round of fluorescence-activated cell sorting, specifically isolating the cells with the highest fluorescent signal. Post-selection analysis of the heterogeneous population via flow cytometry revealed that the proportion of cells with detectable signal had increased from 4% to 25%. Subsequent next-generation sequencing identified a high degree of convergence towards a specific mutation, H81W, which likely serves as the key driver of improved function in this initial round of engineering (Figure 2C).

Building upon the success of our initial round of engineering, we proceeded with a second iteration of mutagenesis, this time shifting our focus further upstream on the sDBM to positions 80, 82, 83, 85, and 86 while stabilizing positions that demonstrated high convergence in the previous round. Specifically, we fixed residues 79 to arginine, 81 to tryptophan, and 84 to proline. We performed MACS and FACS enrichment and selection as described in the preceding round to isolate variants with further enhanced function. Strikingly, post-sort characterization of the heterogeneous population via flow cytometry revealed a substantial increase in the proportion of cells with detectable signal, jumping from roughly 25% to 56%. Subsequent NGS analysis once again demonstrated strong convergence at a specific position that may be responsible for the enhanced RNA interaction. In particular, a K86Y mutation exhibited by far the strongest convergence and is likely the primary driver of improved activity in this second round of engineering (Figure 2C).

### Directed Evolution Yields Engineered Variants with Enhanced Properties

Having successfully enhanced the activity of our enzyme on the surface of yeast by tenfold, we sought to characterize performance *in vitro* to determine if the engineered variants are suitable for application as RNA-interacting HUH-tags. Using the sequencing data from each round of engineering, we identified the most highly enriched variants from rounds one and two, which we designated as E1 and E2, respectively. We then recombinantly expressed and purified these enzymes to test their ability to cleave and covalently link to both DNA and RNA substrates *in vitro*. To better differentiate the function of WT and engineered variants, we implemented more stringent reaction conditions. We reduced the substrate concentration from 30 µM (10X Substrate:Enzyme ratio) to 15 µM (5X ratio). Additionally, we decreased the MnCl_2_ concentration from 1 mM to 50 µM. Finally, we shortened the reaction time from overnight to 30 minutes. Remarkably, while ssDNA covalent linkage remained roughly consistent across the WT, E1, and E2 enzymes, activity on the non-cognate RNA substrate improved dramatically, going from undetectable levels in the WT enzyme under restricted conditions to roughly 60% and 80% in E1 and E2, respectively (Figure 3).

**Figure 3:**
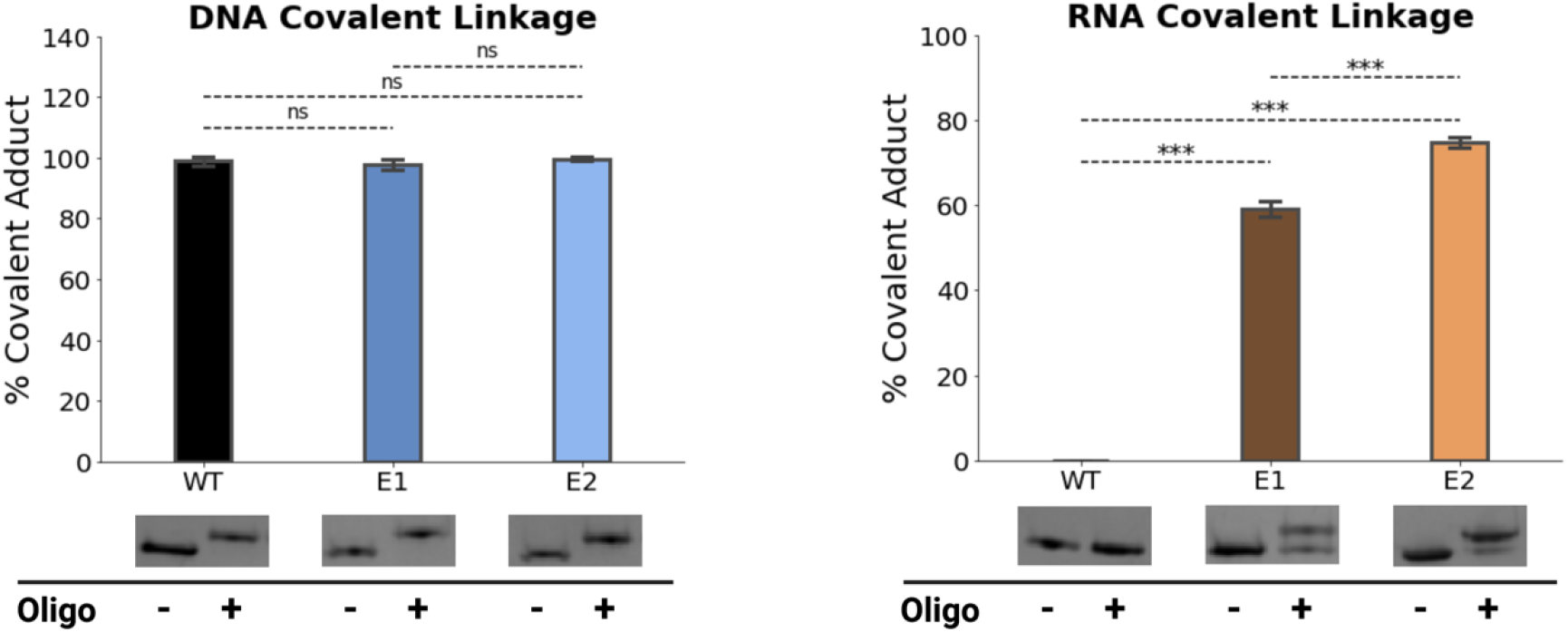
Improved protein-RNA covalent linkage in engineered reps. *in vitro* HUH-tag bioconjugation reactions visualized via SDS-PAGE in triplicate represented as bar plots for both DNA (left) and RNA (right) substrates with the WT, E1, and E2 enzyme variants. Reactions were performed in final concentrations of 3 μM HUH-tag and 15 uM nucleic acid substrate in 50 mM HEPES pH 8.0, 50 mM NaCl, 1 mM DTT, and 50 μM MnCl2 for thirty minutes at 37°C.

These encouraging findings prompted us to investigate the cleavage reaction of the WT and engineered enzymes over time in order to evaluate the extent of the kinetic improvements. To evaluate changes in reaction kinetics, we modified a previously described^39^ continuous molecular beacon cleavage assay that employs single-stranded nucleic acid oligos harboring the enzyme’s specific cleavage motif flanked by a FAM fluorophore and Iowa Black quencher on the 3′ and 5′ ends, respectively. This is a plate reader-based assay where cleavage is read out by an increase in fluorescence upon cleavage of the dual-labeled nucleic acid substrate. Unexpectedly, we found that the efficiency of reactions on DNA substrates improved so dramatically that we had to reduce the reaction efficiency by adding 500 mM NaCl in order to properly observe the kinetics. The engineered variants exhibited dramatically faster reaction rates than their WT counterpart, which was neither anticipated nor detectable in the single time-point covalent linkage analysis (Figure 4). As expected, the reaction efficiency on RNA substrates also improved substantially through successive iterations, with cleavage enhancing from near undetectable levels in the WT enzyme to robust reactivity in the subsequent engineered iterations as seen in the covalent linkage assays (Figure 4).

**Figure 4:**
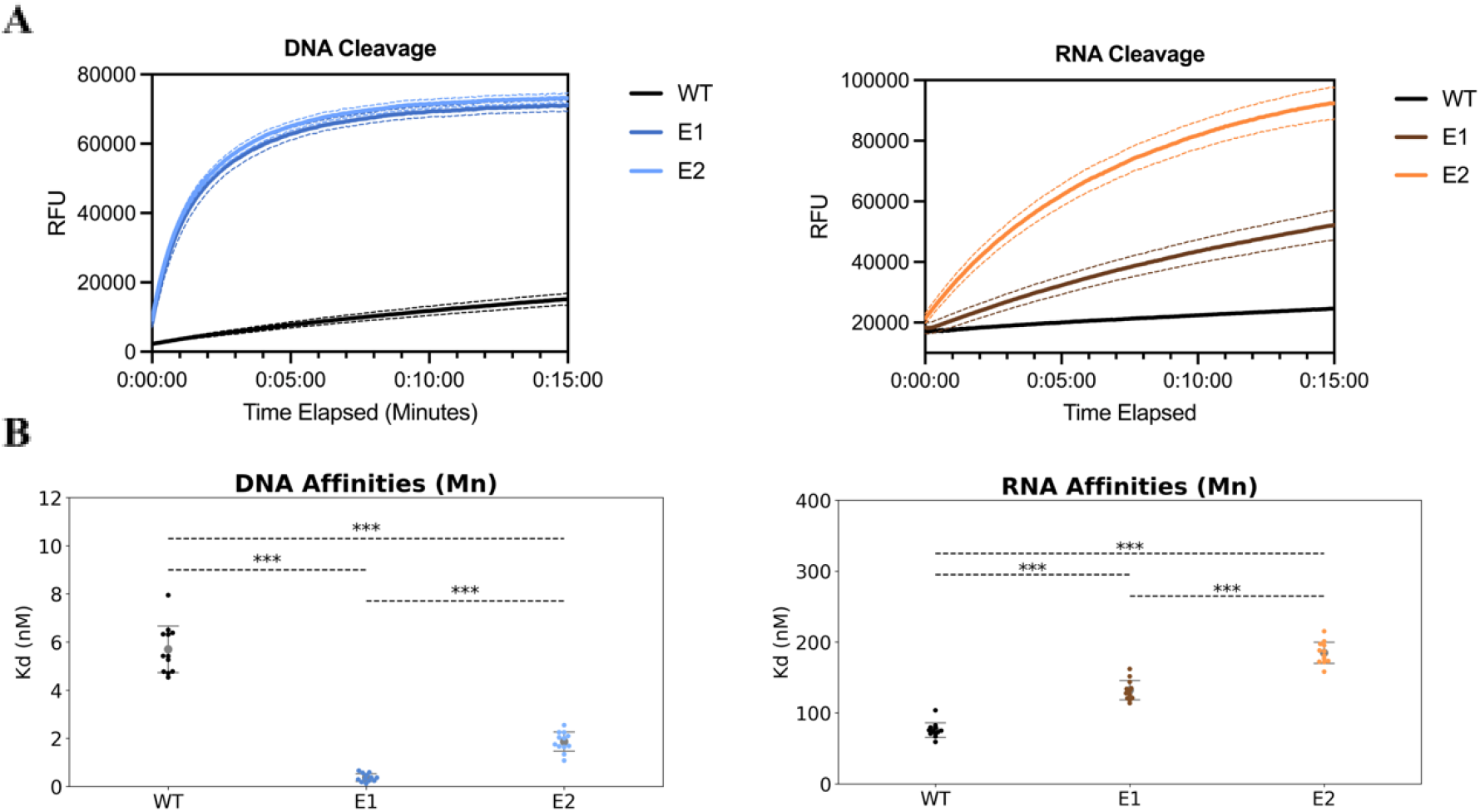
Comparing the kinetics and binding affinities of WT and engineered enzymes across substrates. Continuous molecular beacon cleavage assays (A) and fluorescence polarization substrate binding assays (B) of DNA (left) and RNA (right) for the WT, E1, and E2 versions of PCV2. Molecular beacon cleavage assays were performed with nucleic acid substrates harboring the PCV2 cleavage site with a 5’ quencher and a 3’ fluorophore. All cleavage reactions were performed in a cleavage buffer of either 50 mM NaCl (for RNA cleavage) or 500 mM NaCl (for DNA cleavage), 50 mM HEPES, pH 8.0, 0.05% Tween-20, 1 mM DTT, and 50 µM MnCl_2_. N-terminal SUMO-tagged Rep variants were diluted with the cleavage buffer to 2 µM and the Q/F substrate was diluted to 200 nM in an identical buffer. All fluorescence polarization assays were performed with N-terminal SUMO-tagged catalytically inactive Reps by titration via 1:2 serial dilution starting at 1 µM into a 10 nM solution of a FAM-labeled substrate in a binding buffer of 50 mM NaCl, 50 mM HEPES, pH 8.0, 0.05% Tween-20, 1 mM DTT, and 5 µM MnCl_2_.

We were surprised by the substantial shift in kinetics observed with cognate ssDNA substrates considering that this is not what we were intentionally selecting for. Intrigued by these unexpected results, we sought to better understand the consequences of the changes that we were making to these enzymes throughout the engineering process. To gain a more comprehensive understanding of substrate interaction, we employed a previously described^39^ fluorescence polarization-based assay to quantify changes in affinity across our wild-type and engineered enzymes for both cognate ssDNA and non-cognate ssRNA substrates. Interestingly, although the directed evolution efforts were aimed at enhancing catalysis on RNA, we observed a modest improvement in DNA binding affinity throughout the engineering process. The dissociation constant (Kd) decreased from approximately 6 nM in the WT enzyme to just under 1 nM in the E1 variant, ultimately stabilizing at nearly 2 nM in the E2 variant in the presence of Mn^2+^ as observed by fluorescence polarization. Counterintuitively, the affinity for RNA decreased as the engineering progressed, with the Kd increasing from approximately 90 nM in the WT enzyme to roughly 150 nM in the E1 variant, and finally reaching just under 200 nM in the E2 variant. Taken together, the directed evolution process yielded substantially improved variants with enhanced catalytic activity on RNA substrates as intended. However, these improvements were accompanied by some unanticipated side effects, such as increases to DNA binding affinity and reactivity as well as decreased RNA binding affinity in engineered enzymes. These findings suggest that the mechanism underlying RNA interaction in Reps is likely more complex than initially anticipated, with mutations that enhance catalysis on RNA substrates potentially affecting other properties of the enzyme in unexpected ways. Further investigation into the structural basis of these changes could provide valuable insights into the intricate nature of Rep-RNA interactions and guide future engineering efforts.

### Structure Comparison and in vitro Characterization Confirms Key Mutations

All mutations made throughout the engineering process were clustered in the sDBM, with E1 and E2 differing from the WT enzyme by five and six mutations, respectively (Figure 5A). Mutagenic libraries were designed to modify key positions that make direct and indirect contacts with the ssDNA substrate in an attempt to remodel the active site and enable RNA interaction. In examination of the co-crystal structure of WT PCV2 in complex with its cognate ssDNA substrate^20^, the effects of the key mutations revealed by sequencing can be considered in their local structural contexts. The H81W mutation likely improves the contact made between the enzyme and the +1 A, whereas the role of the K86Y mutation is seemingly less straightforward to interpret (Figure 5A). Interestingly, the engineered variants are strongly enriched for specifically tyrosine at position 86, never phenylalanine, indicating the importance of the hydroxyl group, possibly in mediating a hydrogen bond in an unanticipated location (Figure 4C).

**Figure 5:**
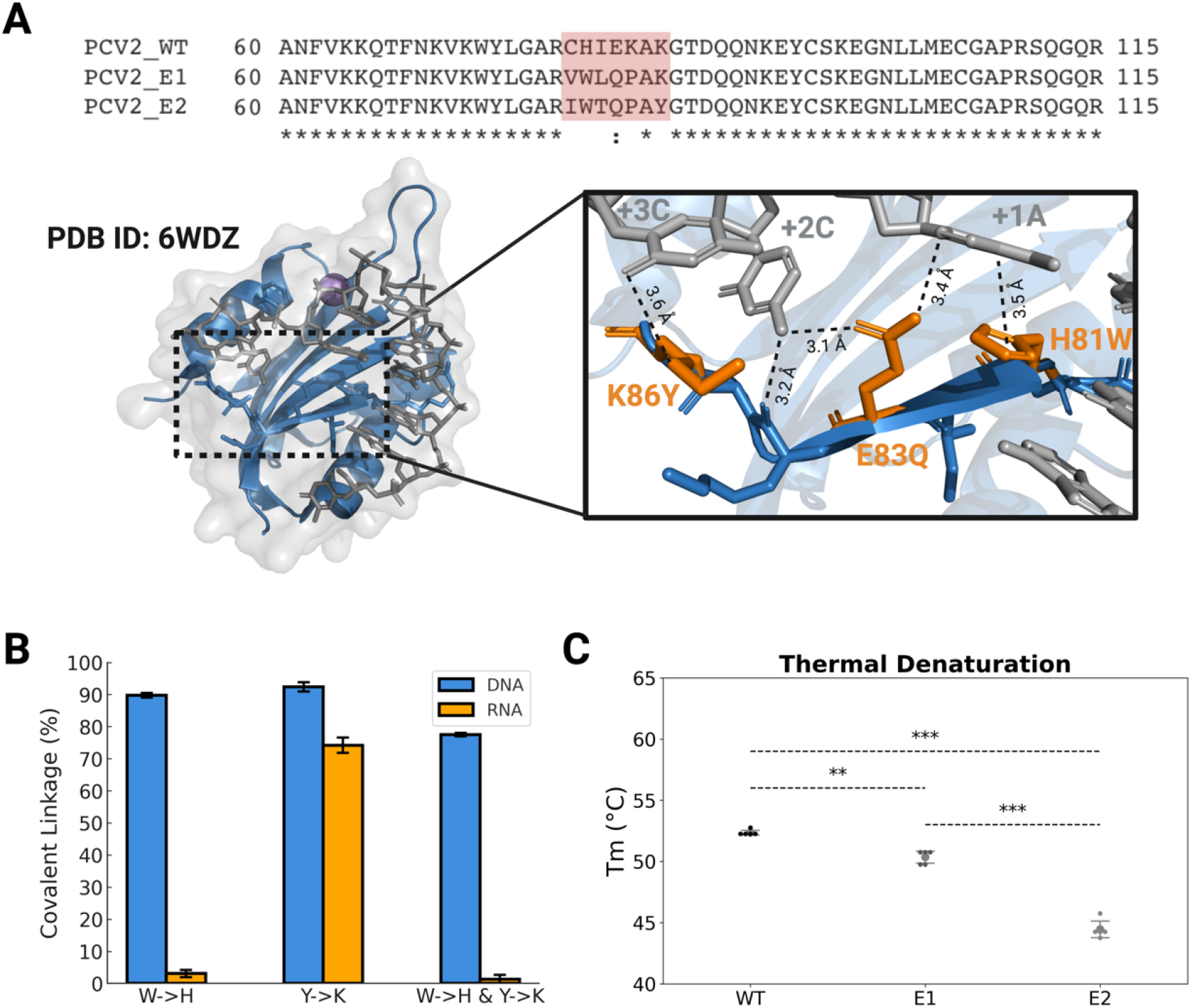
Confirmation of key residues through in vitro reaction. (A) Partial sequence alignment showing the differences between the wildtype and engineered variants above a co-crystal structure of PCV2 in complex with its cognate DNA substrate. The image of the structure highlights the single-stranded DNA bridging motif, which was the primary target of engineering efforts. (B) in vitro HUH-tag bioconjugation reactions visualized via SDS-PAGE in triplicate represented as bar plots for both DNA (left) and RNA (right) substrates with the indicated modified enzyme variants. Reactions were performed in final concentrations of 3 μM HUH-tag and 15 uM substrate in 50 mM HEPES pH 8.0, 50 mM NaCl, 1 mM DTT, and 50 μM MnCl2 for thirty minutes at 37°C. (C) Strip plot demonstrating the differences in Tm as calculated by differential scanning fluorimetry (DSF) after each step of engineering.

To confirm the importance of these two mutations, we reverted them back to their WT residues in context of the E2 enzyme and performed in vitro reactions on DNA and RNA substrates. The reversion of these two mutants had catastrophic effects on enzyme activity, highlighting their crucial role in the enhanced performance of the engineered variants. Specifically, the W81H mutation in E2 nearly ablated RNA interaction altogether under restricted reaction conditions, whereas the Y86K mutation only modestly decreased activity compared to E2 (∼80% linkage to ∼75% in the point mutant). The combination of these two mutations brought RNA interaction to a nearly undetectable level, mirroring the behavior of the WT enzyme (Figure 5B).

Moreover, another unanticipated consequence of the engineering process was the loss of thermal stability. Using differential scanning fluorimetry on WT and engineered versions of our enzymes, we found that the melting temperature (Tm) decreased from roughly 53°C in the WT enzyme to approximately 50°C in E1 and further down to just under 45°C in E2, suggesting that the K86Y mutation may be destabilizing the enzyme. Loss of stability is not uncommon in protein engineering endeavors and is frequently observed in meganucleases engineered using yeast surface display based directed evolution^40^. However, this finding further underscores the complex interplay between the introduced mutations and the overall protein structure and function and how even modest seeming mutations can have unanticipated results.

## Discussion

HUH-tags have emerged as a highly versatile tool for mediating sequence-directed protein-ssDNA covalent bioconjugation, finding utility as integral components of a wide range of application both in vitro and in vivo^20,22-29,41,42^. In this work, we demonstrate that the substrate specificity of HUH-endonucleases can be expanded to enable robust sequence-directed covalent linkage to RNA through directed evolution of a promiscuous rep using yeast surface display. The successful engineering of RNA-interacting HUH-tags not only introduces a versatile new tool for protein-RNA bioconjugation, but also highlights the amenability of these enzymes to engineering of enhanced or novel properties, which could prove useful in other contexts. For example, engineering HUH-endonucleases to improve ssDNA linkage within mammalian cells could potentially lead to enhanced substrate recruitment and thus better outcomes in the emerging ‘click editing’ genome engineering system^42^. Indeed, HUH-tags play a critical role in this novel alternative to prime editing in that they covalently tether a ssDNA template which encodes user-defined mutations to a Cas9-Klenow fragment fusion for site-specific modification of genetic information. Moreover, the ability to engineer HUH-tags with designer sequence specificity would dramatically improve the potential for multiplexing with this bioconjugation platform. While we have identified high efficiency non-cross reactive substrates across pairs, triplets, and even quadruplets of HUH-tags, the vast majority of the sequence space encoded by their substrate length is completely inaccessible to these enzymes – particularly sequences enriched in G content^20,30^. The results presented here lay the foundation of exciting new possibilities for expanding and fine-tuning the functionality of HUH-tags to suit an even broader range of applications in biotechnology and synthetic biology.

While directed evolution is a powerful tool for engineering enzymes with enhanced or novel functions, it is not without limitations. In our engineering of RNA-interacting HUH-tags, we observed that even seemingly comparable amino acid substitutions can lead to a reduction in protein stability. This is a common challenge in protein engineering, as directed evolution tends to prioritize mutations that improve function or fitness without necessarily making compensatory mutations to maintain stability^40,43^. In contrast, natural evolution is able to preserve these relationships more intimately over time when evolving new functions^43^. One potential strategy to mitigate stability loss during engineering is to actively select for higher stability at various stages throughout the process or to computationally engineer greater stability into the protein prior to directed evolution using stabilizing tools like Protein Repair One Stop Shop (PROSS)^44^, FireProt^45^, or ProteinMPNN^46^. We also note that our engineered enzymes exhibited hyperactivity on cognate ssDNA substrates, which could potentially have deleterious effects on sequence specificity. Future engineering efforts could overcome this limitation by focusing on maintaining reactivity while modifying the reaction for designer purposes, such as enhanced activity on RNA or novel sequence specificity. However, given that these nucleases primarily recognize substrates through an indirect shape-based mechanism with minimal contacts to the backbone or sugar groups^20^, it may prove challenging to completely alter or swap substrate specificity from DNA to RNA.

Considering the high structural similarity between ssDNA and ssRNA, it is interesting that only a subset of rep enzymes are capable of promiscuous protein-RNA linkage. Intriguingly, Rep proteins share strong structural similarity with viral RNA recognition motifs^14,47^, suggesting that there may have been an ancient evolutionary ancestor common to DNA and RNA viruses. Interestingly, recombination between co-infecting ssDNA and ssRNA viruses is thought to have given rise to *Cruciviridae*, a family of circular rep-encoding ssDNA (CRESS-DNA) viruses that encode standard rep proteins alongside capsid proteins with high homology to those found in the RNA genome viruses of Tombusviridae^48-50^. Notably, the rep proteins of *Cruciviridae* are most similar to those of *Circoviridae* and bear little resemblance to reps that mediate replication in extrachromosomal plasmids which were thought to have given rise to these viruses^48,49^. The fact that the Reps exhibiting promiscuous RNA-interaction are most closely related to those found in a family of viruses thought to have arisen from recombination between DNA and RNA viruses, is intriguing. This similarity may suggest a more intimate evolutionary relationship between *Cruciviridae* and RNA viruses than previously appreciated. While further investigation is needed to reveal the precise evolutionary implications of this promiscuous activity, these findings highlight the potential significance of the RNA-interacting capabilities observed in circoviruses and provide a new perspective on the evolution of ssDNA viruses.

Protein-RNA covalent linkage is not novel to nature, with most examples originating from viruses. In polioviruses, and other related RNA viruses, the 5’ end of the viral genome is capped with a viral factor referred to as “viral protein genome-linked”, or VPg^51^. This linkage is mediated through a phosphotyrosine bond, similar to the engineered HUH-tag described in this study, and this protein factor is thought to link to RNA and subsequently serve as a primer for genome replication in these viruses^51^. Moreover, a recent study revealed that some phage ADP-ribosyltransferases, which typically link ADP to bacterial proteins during infection, can also accept NAD-capped RNAs as a substrate^52^. This enables the covalent attachment of RNA to host proteins, a process the authors termed “RNAylation”^52^. Even translation relies on the covalent attachment of amino acids to their respective tRNAs, although this is more accurately described as an amino acid-RNA linkage rather than a protein-RNA linkage.

In this work, we have developed a platform that enables robust sequence-directed protein-RNA covalent linkage. By employing yeast surface display to engineer a promiscuous rep protein PCV2, we were able to dramatically improve the efficiency of covalent RNA-protein linkage, achieving a significant enhancement in comparison to the wildtype enzyme. The successful engineering of these RNA-interacting HUH-tags not only provides a versatile tool for covalent protein-RNA bioconjugation but also highlights the potential for further optimization and customization of HUH-tags to suit a wide range of applications. The ability to engineer enhanced or novel activities into these simple and compact fusion partners opens up exciting possibilities for their use in a diversity of applications in biotechnology and synthetic biology.

## Methods

### Design and Amplification of Mutagenic Libraries

Rep-RNA interaction engineering libraries were designed for site saturation mutagenesis of residues in proximity to ssDNA in co-crystal structures, starting with the single-stranded DNA bridging motif (sDBM), which is known to impart sequence specificity in Reps. A maximum of six residues were mutagenized with ‘NNK’ codons (which include all 20 amino acid possibilities plus a single stop codon) in a single library, producing a theoretical library size of 21^6^, or roughly 85.8 million unique amino acid sequences. Alternative ambiguous codons, such as ‘DBK’, were occasionally used to limit residue variation to a subset of amino acids defined from the results of previous engineering stages. Residues showing strong convergence to a single amino acid at a given position would typically be excluded from further rounds of mutagenesis. Further library design was guided by in vitro testing of a panel of the most enriched variants from a previous round of selection when there was no strong convergence to specific residues or characteristics.

Mutagenic libraries were produced in a similar fashion as described by Lambert et al.^40^. Briefly, insert libraries were designed with ∼ 100 base pair overlap with the 5′ and 3′ cleaved ends of the pETcon yeast surface display vector. Assembly PCR was used to generate the mutagenic inserts, which produces a long double-stranded DNA sequence from a collection of short overlapping top- and bottom-strand oligonucleotides. The collection of oligonucleotides used for the assembly reaction was designed by the DNAWorks web server. Site saturation mutagenesis at the desired positions was achieved through the introduction of longer mutagenic oligonucleotide ultramers (IDT) in place of the shorter oligonucleotides spanning a region of interest.

Assembly inserts were produced through two sequential PCR reactions with 2X CloneAmp HiFi PCR premix (Takara). Both reactions were performed in 25 μl volumes, with the first reaction consisting of 1 µL of a 1 µM mixture of the coding region-spanning assembly oligonucleotides and the 2X mastermix, and the second reaction consisting of 2.5 µL each of long forward- and reverse-primers – both with ∼ 100 base pairs of homology to the pETcon vector – and 2.5 µL of the first reaction (non-purified) as a template for amplification. The second reaction was then separated by electrophoresis on a 1% agarose gel and purified via gel extraction. The purified mutagenic insert was then further amplified through numerous additional PCR reactions using the same forward- and reverse primers and purified via either PCR cleanup or gel extraction. Purified mutagenic insert products were pooled and stored at -20°C for future use.

### Media Preparation

2X YPAD rich growth media; recipe for one liter: 20 g Bacto Yeast Extract (ThermoFisher), 40 g Bacto Peptone (Fisher Scientific), 40 g D-glucose (Fisher Scientific), and 100 mg of Adenine hemisulfate (Sigma-Aldrich) dissolved in 990 mL of water, brought to a pH of 6.0 and autoclaved. Once cool, 10 mL of 100X pen/strep antibiotic solution (ThermoFisher) and 500 uL of kanamycin at 50 mg/mL (FisherScientific) was added.

### Yeast Transformation, Culture, and Induction of Surface Expression

The amplified mutagenic libraries were assembled into the pETcon plasmid using the yeast’s natural ability to perform homologous recombination by transformation with a mixture containing cleaved vectors and inserts with ∼ 100 base pairs of homology to its 5′ and 3′ ends. Mutagenic libraries were transformed into EBY100 Saccharomyces cerevisiae (ATCC) using a scaled-up version of the lithium acetate method. Briefly, the night prior to a transformation a small frozen aliquot of non-transformed yeast was cultured in 25 mL of nonselective growth media (2X YPAD). The next morning, this overnight culture was used to inoculate 100 mL of fresh 2X YPAD in a 500 mL baffled shake flask at 30°C shaking at 250 RPM until the culture reached a density of around 100 million cells/mL. 2.5 billion cells (∼ 25 mL) were then transferred to a 50 mL conical tube, spun down for five minutes at 3.5K G, and then resuspended in sterile water. Resuspended cultures were spun down a second time and resuspended in a transformation mixture composed of 2.4 mL 50% PEG 3,350 (Sigma-Aldrich), 360 µL 1M Lithium Acetate (Sigma-Aldrich), 500 uL of denatured salmon sperm DNA (Sigma-Aldrich), and a 340 uL mixture of ∼ 20 µg of the cleaved pETcon yeast surface display plasmid and ∼ 40 µg of mutagenic insert. Cells resuspended in the transformation mixture were then transferred to a 15 mL culture tube and incubated in a water bath at 42°C for 35 minutes, gently mixing every five minutes. Post heatshock, the culture was transferred to a 50 mL conical containing a 25 mL 50:50 mixture of selective media and 20% w/v glucose solution for recovery. Following this, the recovery mixture was added to 500 mL of selective media + glucose in a 2L baffled shake flask and shaken at 250 RPM at 30°C overnight.

The following morning, yeast cells equal to roughly 50X the theoretical diversity of the library were washed in water and passaged into fresh selective media + raffinose and shaken at 250 RPM at 30°C for culture prior to induction. Once the culture reaches a density of roughly 90 - 120 million cells per mL, the desired number of cells are again washed in water and transferred to selective media + galactose at a density of roughly 30 million cells/mL and left at room temperature overnight for induction of cell surface display.

### Protein Engineering

Induced yeast cultures surface-expressing mutagenic libraries were sorted via two rounds of magnet-activated cell sorting (MACS) with biotinylated substrates and magnetic streptavidin beads for functional enrichment, and a final round of fluorescence-activated cell sorting (FACS) with AF647-labeled substrates gated to sort out highly functional variants. The nucleic acid substrates used for selection were RNA versions of simplified CRESS-DNA viral origins of replication (Ori) with 3 to 4 DNA bases appended to the 5′ and 3′ ends (IDT) in order to enhance the stability of an otherwise unmodified RNA substrate.

Briefly, for MACS selection, 1.5 billion induced yeast cells were washed two times with a washing buffer (500 mM KCl, 10 mM NaCl, 50 mM HEPES, pH 7.5) and then were resuspended at 100 million cells per mL in a reaction buffer (50 mM NaCl, 50 mM HEPES, pH 8.0, 1 mM MnCl2, and variable concentration of a 3’ biotinylated selection substrate) for subsequent incubation and reaction at 37°C. Post-incubation, cells were washed twice in washing buffer and resuspended at 100 million cells per mL in a bead-binding buffer (20 mM Tris, pH 7.5, 500 mM NaCl, 1 mM EDTA) and 1 mg of hydrophilic streptavidin magnetic beads (NEB) equilibrated into the same buffer. Yeast-bead solutions were rocked at 4°C for two hours before selection with a magnetic rack by pulling down the magnetic beads and removing the non-bound yeast culture supernatant. Magnetic beads were washed twice with bead-binding buffer and resuspended in 5 mL of selective media with glucose for subsequent growth for 16 hours. Following this, the cultures were spun down and resuspended with 1 mL of selective media with raffinose, incubated on a magnetic rack at room temperature for five minutes, and the non-bound supernatant was transferred to fresh selective media with raffinose for further growth and induction. The selection substrate was titrated down throughout the engineering process, reactions were performed with a 100 nM concentration for the first library, 50 nM for the second, and so on. Additionally, the first round of magnet selection on a given library was reacted with substrate for 15 minutes for functional enrichment and the second round for 5 minutes to further select for highly functional variants.

Briefly, for FACS selection, the desired number of induced yeast cells (typically ∼ 100 million) were washed two times with the washing buffer and were then resuspended at 100 million/mL in a reaction buffer (50 mM NaCl, 50 mM HEPES, pH 8.0, 500 uM MnCl_2_ and 50 nM AF647-labeled substrate) for subsequent incubation and reaction at 37°C. Post-incubation, cells were washed twice in washing buffer and resuspended at 100 million cells/mL in a staining buffer (200 mM KCl, 10 mM NaCl, 50 mM HEPES, pH 7.5, 1:100 dilution of a FITC-labeled anti-Myc antibody (ICL)) and then rocked in the cold room for up to two hours. Following this, cells were spun down and resuspended in this same staining buffer at roughly 50 million cells/mL and held on ice until sorted on a BD FACSAria II P0287 cell sorter at the University of Minnesota flow cytometry resource. Flow sorted cells were then transferred to fresh selective media + glucose and allowed to shake overnight at 250 RPM at 30°C prior to further culture of manipulation.

### Flow Cytometry Analysis of Expression and Activity on the Yeast Surface

Enzymes displayed on the yeast surface were analyzed for activity and expression in a manner similar to that of FACS selection, but on a much smaller scale. Stained cells were analyzed on a BD Accuri benchtop flow cytometer (BD) using the FITC and AF647 channels with no compensation necessary.

### Molecular Cloning

The WT 6xHis-SUMO-Rep PCV2 plasmid was generated in a previous study^20^. Briefly, a WT Rep sequence was cloned into the pTD68/6xHis-SUMO expression vector as a geneblock (IDT) with the In-Fusion HD Cloning Kit (Takara) using BamHI and XhoI restriction sites (NEB). Engineered variant plasmids for protein expression were either generated in an identical manner or through site-directed mutagenesis using the same cloning kit per manufacturer instructions. Enzymes containing point mutations were also created via site-directed mutagenesis as described above. Each plasmid was sequence confirmed with Sanger sequencing (Genewiz) prior to use.

### Protein Expression and Purification

Rep constructs were expressed in BL21(DE3) competent E. coli cells (Agilent) in a 1 L volume with LB broth. Temperature was reduced from 37°C to 18°C after the culture OD600 reached 0.8 and cells were induced with 0.5 mM IPTG (Sigma Aldrich) and incubated overnight. Cells were lysed in a lysis buffer (250 mM NaCl, 50 mM HEPES, pH 8.0, 1 mM EDTA) with a complete protease inhibitor tablet (Pierce) and pulse sonicated at 4°C. Clarified supernatant was batch bound with Ni-NTA HisPur agarose beads (ThermoFisher) for 1 hour, loaded onto a gravity column, washed with 30 column volumes of wash buffer (250 mM NaCl, 50 mM HEPES, pH 8.0, 1 mM EDTA, 30 mM imidazole), and finally eluted with elution buffer (250 mM NaCl, 50 mM HEPES, pH 8.0, 1 mM EDTA, 300 mM imidazole). Purification was assessed via SDS-PAGE gel analysis and protein-containing fractions were dialyzed (250 mM NaCl, 50 mM HEPES, pH 8.0, 1 mM EDTA, 1 mM DTT) overnight at 4°C. Protein was further purified and buffer exchanged using the Superdex 300 Increase 10/300 GL (GE Healthcare) size exclusion column into the lysis buffer for storage and characterization. For the production of SUMO-free protein, approximately 30 µg 6xHis-ULP1 (SUMO-specific protease) per liter of culture was added to the protein samples prior to overnight dialysis. Following this, protein samples were then batch-bound a second time with Ni-NTA HisPur agarose beads to remove cleaved 6xHis-SUMO and 6xHis-ULP1 for subsequent buffer exchange as described above.

### In vitro HUH Cleavage Reactions

In vitro oligonucleotide cleavage reactions were carried out using final concentrations of 3 µM N-terminal SUMO-tagged Rep variant and 30 µM single-stranded nucleic acid Ori substrate (IDT) in a cleavage buffer of 50 mM NaCl, 50 mM HEPES, pH 8.0, 1 mM DTT, and 1 mM MnCl_2_ and incubated overnight. Alternatively, restrictive reactions were carried out using final concentrations of 3 µM N-terminal SUMO-tagged Rep variant and 15 µM single-stranded nucleic acid Ori substrate in a cleavage buffer of 50 mM NaCl, 50 mM HEPES, pH 8.0, 1 mM DTT, and 50 µM MnCl_2_ and incubated for 30 minutes. Reactions were incubated at 37°C before subsequent denaturation and SDS-PAGE analysis.

### Fluorescence Anisotropy Assays

The affinity of engineered and WT Reps for ssDNA and ssRNA were measured via fluorescence anisotropy assays as previously described^39^. 3’ fluorescein (FAM) labeled nucleic acid Ori substrates were used in all anisotropy experiments (IDT). N-terminal SUMO-tagged catalytically inactive Reps were titrated via 1:2 serial dilution starting at 1 µM into a 10 nM solution of a FAM-labeled substrate in a binding buffer of 50 mM NaCl, 50 mM HEPES, pH 8.0, 0.05% Tween-20, 1 mM DTT, and 5 µM MnCl_2_. A single binding reaction was assembled in quadruplicate using four rows across a 96-well plate and three independent replicates of this reaction were performed across separate days. Reactions were incubated for 30 minutes at room temperature to ensure equilibrium and centrifuged at 2,000 G for two minutes before measurement. Mean dissociation constants (K_D_) were calculated across three separate reactions (n=12) using the equations described below. In order to calculate anisotropy from parallel and perpendicular fluorescence intensity values, data were fit to the following equation:

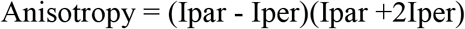

Where Ipar equals parallel fluorescence intensity and Iper equals perpendicular fluorescence intensity. In order to calculate K_D_ under non-saturating ligand conditions, data is fit to a quadratic model:

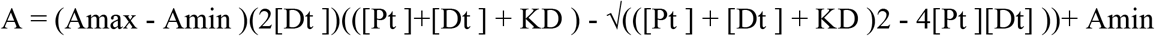

Where [Dt] is the total amount of labeled ssDNA, [Pt] is the total amount of Rep protein, K_D_ is the dissociation constant, Amax is the anisotropy ceiling, and Amin is the anisotropy floor. All anisotropy data were collected using a Synergy Neo2 Hybrid Multi-Mode Plate Reader and the Green FP Filter Cube (Agilent) with an excitation wavelength of 485 nm (20 nm bandwidth). Emissions of 528 nm were measured at ambient temperature. Measurements were fit and plotted in GraphPad Prism according to the above equations.

### Molecular Beacon Cleavage Assays

The cleavage rate of engineered and WT Rep variants was evaluated with a molecular beacon assay using either a ssDNA or ssRNA probe harboring an Ori-derived sequence labeled with a 5’ IowaBlack-FQ quencher and a 3’ FAM (Q/F, IDT). All cleavage reactions were performed in a cleavage buffer of either 50 mM NaCl (for RNA cleavage) or 500 mM NaCl (for DNA cleavage), 50 mM HEPES, pH 8.0, 0.05% Tween-20, 1 mM DTT, and 50 µM MnCl_2_. N-terminal SUMO-tagged Rep variants were diluted with the cleavage buffer to 2 µM and the Q/F substrate was diluted to 200 nM in an identical buffer. In a black 96-well plate, 100 µL of the Q/F substrate was added to each well and inserted into a Synergy Neo2 Hybrid Multi-Mode Plate Reader (Agilent). Using the injector module, 100 µL of a Rep variant was added to each well, initiating the reaction. Fluorescent signal was collected by the plate reader using the monochromator function with an excitation wavelength of 485 ± 20 nm and emission collected using a 528 ± 20 nm bandpass filter at room temperature. Emission values for all 8 reactions were collected every 5 seconds for 15 minutes. At a minimum, three independent replicates were collected for each condition.

### Differential Scanning Fluorimetry

DSF experiments were conducted using a CFX96 Touch™ Real-Time PCR Detection System (Bio-Rad) with FRET channel detection. The assay was performed in skirted qPCR plates with a final reaction volume of 35 μL per well. The reaction mixture consisted of 2 μM enzyme, 600 mM NaCl, 20 mM HEPES (pH 8.0), and 10X SYPRO Orange dye (Thermo Fisher). A master mix was prepared for each protein of interest, containing 2X DSF buffer (1.2 M NaCl, 40 mM HEPES pH 8.0), 30 μM protein stock, and water. SYPRO Orange dye was diluted separately to a 70X working solution using 2X DSF buffer and water. The plate was prepared with five replicates each of water-only, buffer-only, and protein samples. Four out of five wells for each condition received 5 μL of 70X SYPRO Orange dye, with one well left dye-free as a control. The plate was then centrifuged at 2000 G for two minutes. Thermal denaturation was measured using a temperature gradient from 20°C to 95°C, with a ramp rate of 1°C/min. Melting temperatures (Tm) were determined from the first derivative of the thermal denaturation profiles using GraphPad PRISM (v. 6.07).

## Author contributions

ATS designed and performed all directed evolution experiments; ATS, NSBD, and AJK, cloned, expressed, and purified enzyme variants; ATS designed and performed all in vitro characterization assays; KJT discovered Rep-RNA promiscuity; ATS prepared and wrote the manuscript; HA acquired funding; WRG acquired funding, aided in experimental design, oversaw data collection and analysis, and wrote the manuscript. All listed authors gave approval of the final version of this manuscript.

